# Exploring functional variation affecting ceRNA regulation in humans

**DOI:** 10.1101/016865

**Authors:** Mulin Jun Li, Jiexing Wu, Peng Jiang, Wei Li, Yun Zhu, Daniel Fernandez, Russell J. H. Ryan, Yiwen Chen, Junwen Wang, Jun S. Liu, X. Shirley Liu

## Abstract

MicroRNA (miRNA) sponges have been shown to function as competing endogenous RNAs (ceRNAs) to regulate the expression of other miRNA targets in the network by sequestering available miRNAs. As the first systematic investigation of the genome-wide genetic effect on ceRNA regulation, we applied multivariate response regression and identified widespread genetic variations that are associated with ceRNA competition using 462 Geuvadis RNA-seq data in multiple human populations. We showed that SNPs in gene 3’UTRs at the miRNA seed binding regions can simultaneously regulate gene expression changes in both *cis* and *trans* by the ceRNA mechanism. We termed these loci as endogenous miRNA sponge expression quantitative trait loci or “emsQTLs”, and found that a large number of them were unexplored in conventional eQTL mapping. We identified many emsQTLs are undergoing recent positive selection in different human populations. Using GWAS results, we found that emsQTLs are significantly enriched in traits/diseases associated loci. Functional prediction and prioritization extend our understanding on causality of emsQTL allele in disease pathways. We illustrated that emsQTL can synchronously regulate the expression of tumor suppressor and oncogene through ceRNA competition in angiogenesis. Together these results provide a distinct catalog and characterization of functional noncoding regulatory variants that control ceRNA crosstalk.

## Introduction

Recent RNA biology has revealed that specific RNAs can operate as ceRNAs to titrate away the pools of active miRNAs, indirectly regulating the expression of other transcripts targeted by the same set of miRNAs. Endogenous miRNA sponges, including mRNA, pseudogene transcripts, long non-coding RNAs and circular RNAs, have been discovered in quick succession^1; 2^, and play critical roles in cellular metabolism and disease development^3; 4^. Besides the different type of transcripts, it has been proposed that structure variations can perturb ceRNA competition and initiate subsequent disease pathway, such as large deletion and insertion, copy number variation, as well as chromosomal translocation^5-7^. ceRNA crosstalk is determined by miRNA response elements (MRE), which encode the ceRNA regulatory network and sustain the dynamic equilibrium for all ceRNAs and miRNAs within the network^8^. Under this circumstance, any genetic event affecting MRE will trigger the perturbation of ceRNA regulation by titrating miRNA availability. One example is SNP rs17228616, which disrupts the interaction between miR-608 and AChE and suppresses other miR-608 targets such as CDC42 and IL6^9^. Nevertheless, there has been no study to our knowledge that systematically tests whether such genetic effect is pervasive in the human evolution and populations.

Quantitative traits, such as gene expression and epigenetic modifications, are thought to be largely heritable during species evolution^10; 11^. Recently next generation sequencing technologies have enabled us to unravel the correlation between genetic variants and different molecular phenotypes, and facilitated the discovery of many human quantitative trait loci (QTLs)^12^. For miRNA-related molecular traits, researchers have discovered many miRNA gene expression QTLs (miRNA-eQTL) that control miRNA gene expression^13-16^ and 3’ untranslated region (3’UTR)-eQTLs that are associated with miRNA target expression^17; 18^. With a better knowledge of ceRNA crosstalk and competition, it is very important to understand how genetic polymorphisms shape human ceRNA regulation. Using 1000 Genomes Project genotype and Geuvadis RNA-seq phenotype data, our study here represents the first effort to investigate the genetic variants associated with ceRNA crosstalk.

## Material and Methods

### The logic for detecting genetic variants affecting ceRNA regulation

We assume that genetic variants, such as SNPs and indels that affect miRNA response elements (MREs) will perturb ceRNA regulation by titrating miRNA availability. Specifically, a causal variation in the seed region of miRNA binding site can introduce different consequences of miRNA-target interaction, by creating or erasing an MRE, or strengthening or weakening an MRE in a ceRNA driver gene (ceD). These subtle changes could perturb the original miRNA and ceRNA regulatory network and the dynamic distribution of other miRNA target genes. Since we focused on the independent effect of one SNP or indel in one MRE of a ceD, the variants will mostly affect the distribution of associated miRNA and its direct targets (ceTs) although the cascade effect could impact other miRNAs and ceRNAs (Supplemental Figure S1). To simplify the investigation of variant effect, we focused on unique miRNA-centered regulatory network in this study. A SNP or indel that creates or strengthens an MRE will decrease the expression of its own host gene (refer to ceD in this study) and increase the expression of the corresponding ceT, while a SNP or indel that erases or weakens an MRE will have the opposite effect. We therefore termed these variants candidate endogenous miRNA sponge (ceRNA) expression quantitative trait loci (emsQTLs).

### Expression data and genotype data

We used Geuvadis RNA sequencing data and small RNA sequencing data of 462 unrelated human lymphoblastoid cell line samples from the CEPH (CEU), Finns (FIN), British (GBR), Toscani (TSI) and Yoruba (YRI) populations in 1000 Genomes project^16; 19^. To be consistent with Geuvadis eQTL detection framework, we directly utilized Geuvadis quantifications for gene and miRNA expression. Geuvadis also provides 462 human genotypes as processed VCFs, of which 421 samples are from 1000 Genomes project Phase1 release v3 and 42 samples are from 1000 Genomes project Phase 2 Omni 2.5M genotype array with imputation. We used major allele frequency (MAF) > 5% as cutoff to select variants for downstream analysis.

### Construct variant-miRNA-ceD-ceT unit

We extracted human 3’UTR sequences according to GENCODE^20^ V12 annotation (consistent with Geuvadis RNA-seq quantification) and 714 Geuvadis quantified miRNA sequences from miRBase^21^. We mapped Geuvadis biallelic genotypes to 3’UTR sequences to construct reference and mutant miRNA targets. Assuming variant independence, we used TargetScan 6.2 to predict miRNA-target relationship^22^. To select reliable miRNA-target pairs, we filtered the prediction by TargetScan Context+ Score (TargetScan cutoff: < -0.310 for 8mer, < -0.161 for 7mer-m8, < -0.099 for 7mer-1A). We also adopted ViennaRNA Package^23^ to estimate the secondary structure-based energies of mRNA with (ΔG_duplex_) or without (ΔG_open_) interaction with miRNA, and used RNAhybrid to display the binding pattern^23^. We then selected those “variant-miRNA-target” units that meet any of the following three criteria: 1) target gain: sequence with alternative allele is a miRNA target site but not for reference allele; 2) target loss: reference allele is but alternative allele is not a miRNA target site; 3) change of context+ score between target sites with reference allele and alternative allele. We treated the miRNA targets meeting the above criteria as putative ceDs. For each selected variant-miRNA-ceD unit, we further searched candidate ceTs under the control of the same miRNA as the ceD (on either reference and mutant 3’UTR) according to TargetScan prediction. Finally, we tested the association between a genetic variant and each ceD-ceT pair under a miRNA-centered regulatory network.

### Control for confounding factors

The quantifications of gene and miRNA expression are usually affected by different technical variations and hidden factors, which will reduce power to interpret expression variability caused by genetic factors. To maximize the emsQTL detection power, we used PEER^24^ to estimate the hidden confounding factors (K) in expression quantifications, and selected the first ten PEER factors (K=10) according to the performance report of Geuvadis eQTL detection. We also calculated expression residuals for miRNA quantifications after accounting for estimated PEER factors, which serve as an essential confounder to control miRNA expression variability. In order to control population stratification, we performed principal component analysis to estimate principal components (PCs) for 462 individual genotype data and selected first three PCs as additional model covariates in the QTL analyses.

### Multivariate linear model

We wanted to test if the variant regulates the expression of ceD and ceT in a reciprocal pattern due to the ceRNA competition. This can normally be achieved by a two-step linear regression on gene expressions against genotype and a set of confounders, with ceD selection in the first step and ceT selection in the second step. However, expressions of the ceD and ceT pairs usually show positive correlation at the functional allele state, suggesting that we could benefit from modeling the relationship between these two responses jointly. Therefore, we used the multivariate linear regression on two responses to simultaneously model the genetic contribution on variability of ceRNA regulation. For each variant-miRNA-ceD-ceT unit, we considered gene expression, measured as the sum of all transcript RPKMs of ceD (*Y*_*d*_) and ceT (*Y*_*t*_), as two dependent variables and transform them to standard normal. We further incorporated the following confounding factors beside the individual genotype (*G*): the PEER residual of miRNA expression (*M*_*r*_), the 10 PEER factors of ceD expression (*PF*_*d*_), the 10 PEER factors of ceT expression (*PF*_*t*_), and first three PCs of individual genotype (*PC*). The separate regression model is shown below:

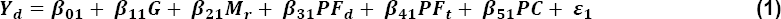

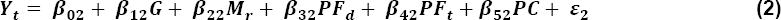

The multivariate regression on both ceD and ceT:

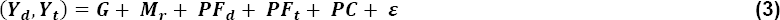

where we simultaneously measured two responses *Y*_*d*_ and *Y*_*t*_, and the same set of predictors on each sample unit. *ε* = (*ε*_1_, *ε*_2_)′ has expectation zero and an unknown covariance matrix. The errors associated with different responses on the same sample unit may have different variances and may be correlated.

Thus, the variant effect for ceD and ceT can be estimated by the corresponding coefficients on genotype (*β*_*d*_ and *β*_*t*_), and we measured this multivariate model by Pillai’s trace test statistics. However, in our definition of emsQTLs, we required opposite expression trend between ceD and each of its ceTs accompanying the genotype change (AA = 0, AB = 1 and BB = 2), so we further filtered variants to only keep those with their *β*_*d*_ and *β*_*t*_ values having opposite signs (*β*_*d*_ × *β*_*t*_ < 0). We finally reported the variant-miRNA-ceD-ceT unit with Benjamini-Hochberg false discovery rate < 0.05.

### emsQTL attributes analysis

To be consistent with the Geuvadis RNA-seq quantification, we utilized GENCODE V12 gene annotation to investigate the functional properties of ceDs and ceTs. We also used DAVID to find enriched gene annotations and pathways^25^. We used SNVrap to annotate genetic variants^26; 27^, and grouped the GWAS traits according to ontology mapping (human phenotype ontology and disease ontology) of GWASdb^28^.

### Evolutionary analysis

We used six statistical measurements, including difference of derived allele frequency (DDAF)^29^, fixation Index (*F*_ST_)^30^, Tajima’s D (TD)^31^, integrated haplotype score (iHS)^32^, cross-population extended haplotype homozygosity (XPEHH)^33^ and cross-population composite likelihood ratio (XPCLR)^34^, to evaluate signals of positive selection on each detected emsQTL SNP using genotype data from five populations of 1000 Genomes project (CEU, FIN, GBR, TSI and YRI). Statistical significance was evaluated by dbPSHP^35^, and we only kept emsQTL SNP with at least one statistically significant score out of the six scores (Supplemental Table S1). Hierarchical clustering was used to cluster the selected emsQTLs according to their derived allele frequencies (DAF).

### Functional prediction of variant effect

We adopted TargetScan context+ score and combined interaction energy score^36^ (ΔΔG = ΔG_duplex_ - ΔG_open_) to measure the alteration of binding affinity for each miRNA-target interaction. We then calculated the distance of context+ score (Δcontext+ score) and the distance of combined interaction energy (ΔΔΔG) between alternative allele and reference allele, which are used to predict the miRNA-target binding affinity change. In analogy with *β*_*d*_, a negative score represents the gain-of-function effect, whereas a positive score represents the loss-of-function effect.

### GWAS enrichment

GWAS traits/diseases associated SNPs (TASs) were collected from GWASdb, NHGRI GWAS Catalog^34^, HuGE^37^, PheGenI^38^ and GRASP^35^, resulting in 33,645 significant SNPs with *P* < 1E-5. To link the signal in the linkage disequilibrium (LD) region, we calculated SNP correlations by MATCH based on the 1000 Genomes project super population for EUR. We obtained all linked SNPs with r^2^ > 0.8 for each GWAS leading TAS and identified emsQTLs overlapping with this expanded list. To test the enrichment of emsQTLs in GWAS signals, we prepared two background datasets for the SNP distribution in the miRNA binding site. We mapped all 1000 Genomes project SNPs into miRNA seed binding region predicted by TargetScan as the first background. We further required that SNPs in first background should have changed binding affinity (Δcontext+ score is not equal to zero) under different alleles to form the second strict background. Using those two backgrounds, we overlapped them with extended GWAS signals and tested the enrichment by hypergeometric test.

## Results

### Genetic Effects on ceRNA Regulation in Human Populations

Our variant selection pipeline (Supplemental Figure S2) successfully mapped 3,544 unique genetic variants (including 3,263 SNPs and 281 indels) on miRNA seed binding sites in the 3’UTR of 2,753 genes (putative ceDs) using 462 Geuvadis individuals. These loci have shown differentiated interaction patterns with 439 miRNAs (out of 714 profiled by Geuvadis) between the reference and alternative alleles. For each putative ceD, we can match over hundreds of other genes (ceTs) which are targeted by the same miRNA according to TargetScan prediction. We applied the multivariate linear regression (in Equation 3) to detect genomic loci that regulate expression levels of a ceD and each of the corresponding ceT. The model includes several essential confounding factors as regressors, including miRNA expression for controlling variability of miRNA concentration among individuals, PEER factors (K) estimating the hidden confounding factors of RNA-seq quantifications, and principal components of individual genotype accounting for population stratification. This multivariate linear model can simultaneously test for two responses of both ceD and ceT expressions and take advantage of the potentially correlated nature between ceD and ceT. After controlling at the FDR of 5%, we further filtered out units with *β*_*d*_ × *β*_*t*_ > 0 and only retained those showing opposite signs of association between genetic variants and gene expression of the two ceRNAs.

#### Genome-wide detection of emsQTLs in different populations

We applied the model and the filtering strategy to five Geuvadis populations independently and successfully detected many emsQTLs. We found 67 (CEU, 91 individuals), 97 (FIN, 95 individuals), 106 (GBR, 94 individuals), 66 (TSI, 93 individuals) and 47 (YRI, 89 individuals) significant associations of unique variant at 5% FDR respectively (Supplemental Table S2). To improve the detection power, we merged the four European subpopulations (EUR, 373 individuals) and detected 387 total significant emsQTLs and 1,875 variant-miRNA-ceD-ceT units (Supplemental Table S3, Figure 1). In the 387 emsQTLs associated with the EUR population, 344 are SNPs and remainings are indels (Figure 2A), suggesting an enrichment of indels over SNPs in affecting miRNA-target interaction and subsequent ceRNA regulation (*P* = 0.04, chi-square test). Different from eQTL detection, emsQTL are not only directly associated with its located ceD in *cis*, but also associated with the corresponding ceT in *trans* through their common miRNA regulator. To investigate if some functional eQTLs can be explained by ceRNA regulation, we checked the number of emsQTLs that overlap with the Geuvadis fine mapping (“the best”) eQTL result from the EUR population, and found to our surprise only 6 overlaps. However, when we considered the Geuvadis all mapped eQTLs, the overlap of emsQTLs is significantly improved (43%, Figure 2B). This suggests that other independent associations may exist in the linked region of each finely mapped eQTL, and the emsQTLs spectrum have pinpointed many additional associations that were missed by conventional eQTL analyses.

**Figure 1:**
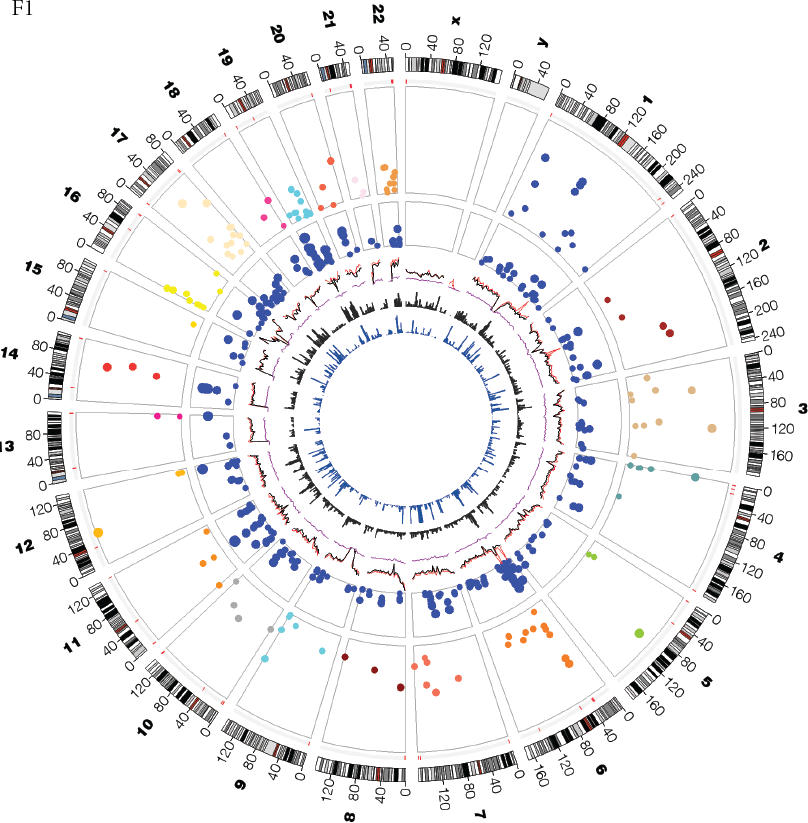
Circos plot of all detected emsQTLs. Features or glyphs are displayed from the outer to the inner, include the number of chromosome, the chromosome ideograms, copy number variation hotspots (red region), Manhattan plot for emsQTLs with –log10(*P*-value), Manhattan plot for GWAS TASs in miRNA binding site predicted by TargetScan, genome variant density (red: dbSNP, black: 1000 Genomes, purple: HapMap 3), OMIM gene distribution and disease-susceptible region distribution.

**Figure 2:**
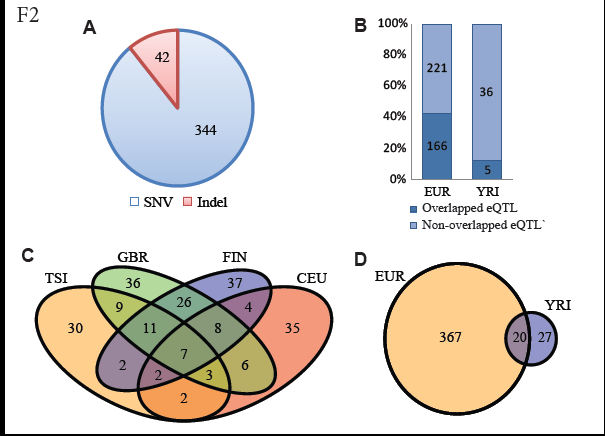
Genome-wide detection of emsQTLs. (A) The proportion of emsQTLs for SNP and indel. (B) The overlapping of emsQTLs in Geuvadis eQTLs. (C) Venn diagram of emsQTLs in European populations. (D) Venn diagram of emsQTL between European and African populations.

#### Positive selection on ceRNA regulation

By simply overlapping the emsQTLs in different subpopulations, we can also find many loci in common or specific to each population (Figure 2C and 2D). Similar patterns can be observed for related miRNAs, ceDs and ceTs of emsQTLs as well (Supplemental Figure S3). Surprisingly, the number of detected emsQTLs is drastically different among different subpopulations despite their similar sample sizes and expected statistical power. When we ranked the number of detected emsQTLs of each subpopulation in ascending order, we found that the sequence of subpopulations (YRI, TSI, CEU, FIN, GBR) follows precisely the human migration path in Europe (Supplemental Figure S4). This phenomenon may suggest that the recent positive selection is shaping the evolution of ceRNA regulation in human populations due to migration and subsequent adaption. To investigate whether emsQTLs are putative targets of the recent positive selection, we screened emsQTLs using six statistical measures (DDAF, *F*_ST_, TD, iHS, XPEHH, and XPCLR) for each subpopulation. We found 46 emsQTLs with positively selected signals for at least one of the measures according to their corresponding empirical thresholds (Supplemental Table S4). Hierarchical clustering for derived allele frequencies of these 46 genetic variants clearly recovers the population relationship and shows distinct pattern on individual locus (Figure 3A). For example, one of YRI-specific emsQTLs rs1050286 (*P* = 8.11E-6) shows disparate derived allele frequency between African (DAF of YRI: 0.89) and European population (DAF of TSI: 0.44; CEU: 0.47; GBR: 0.54; FIN: 0.53) (Supplemental Figure S5). SNP rs1050286 has four measures passing the significant cutoff in the YRI population (DDAF: 0.266; *F*_ST_: 0.12; iHS: 2.285; XPEHH: 1.24), suggesting its likely positive selection in YRI. Long-range haplotype analysis also confirmed the selective sweep around this locus (Figure 3B and 3C). This population-specific emsQTL was detected by our model to regulate ceD *OLR1* and ceT *HORMAD2* by miR-149-5p, and derived allele A enhances the binding affinity for miR-149-5p and *OLR1* according to direction of *β*_*d*_ coefficient (positive value). Therefore, the gene expression of *OLR1* is down-regulated in allele A state, which increases the gene expression of target ceRNA *HORMAD2* from miRNA sponge effect. Previous studies have implicated the overexpression of *OLR1* gene in many diseases including alzheimer’s disease, atherosclerosis, myocardial infarction, obesity, dyslipidemia, and cancer^39-41^. Our ceRNA analysis suggests that the protective role of allele A on rs1050286 in African population arises from miR-149-5p shifting its targets from *HORMAD2* to *OLR1* and reduces the *OLR1* expression through ceRNA competition.

**Figure 3:**
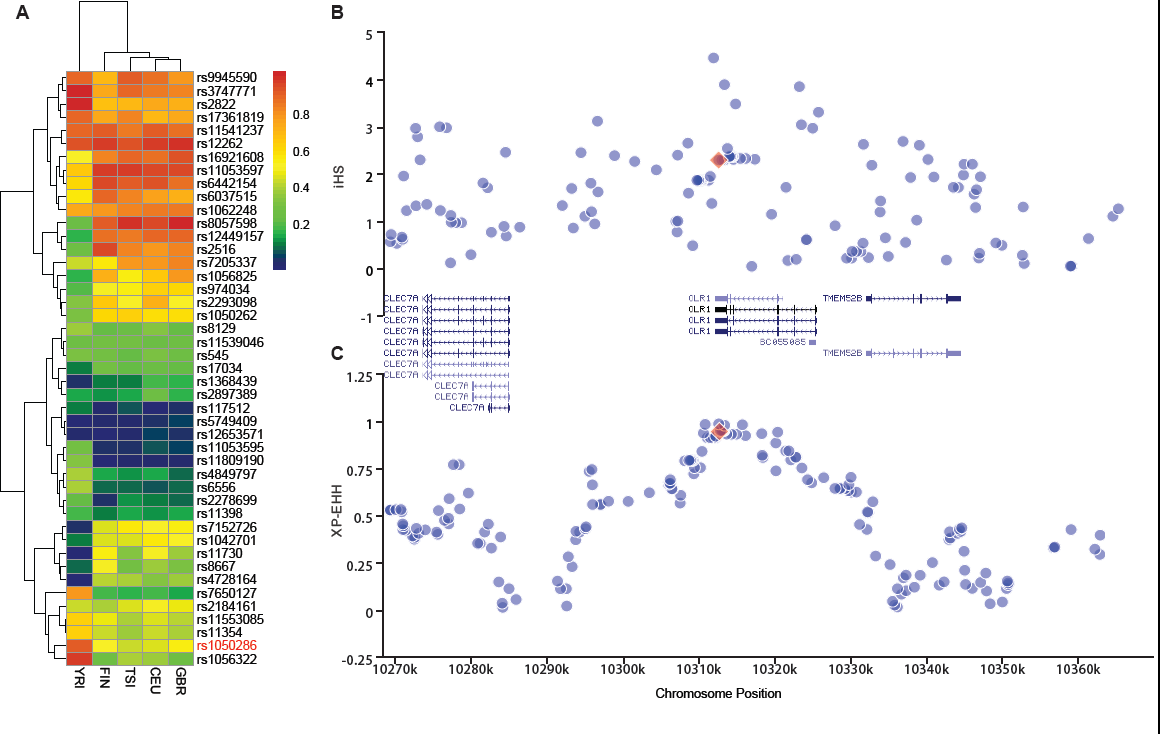
The positive selection of emsQTLs. (A) The hierarchical clustering of according to derived allele frequency for 46 putatively positive selected emsQTLs in different population. (B) iHS scores in rs1050286 locus for YRI. (C) XPEHH scores in rs1050286 locus for YRI.

### Putative Causality of emsQTLs

#### Evaluating emsQTLs properties with functional investigations

Functional interpretation of emsQTLs is pivotal to our understanding their underlying biological mechanisms and phenotype causality. Coefficients of ceD and ceT in our regression model can reflect the degree of gene expression perturbation under different genotypes. Using the EUR 387 emsQTLs, we found a majority of *β*_*d*_ and *β*_*t*_ to be small (< 1) in the 1,875 significant variant-miRNA-ceD-ceT units (Figure 4A and 4B), which indicates a moderate effect of these genetic variations in increasing the precision of target gene expression and ceRNA regulation. To investigate whether the 298 emsQTL-associated ceDs and 1,459 ceTs are engaged in important biological processes, we performed DAVID gene-annotation enrichment analysis for these two gene sets. ceDs were enriched in nucleotides binding, suppressor of cytokine signaling family protein binding and DNA repair (Supplemental Figure S6), while ceTs were enriched in sialyltransferase function, tyrosine protein kinase function, positive gene regulation, transcription regulation and cell proliferation (Supplemental Figure S7). These may indicate that many emsQTLs affect expressions of transcriptional regulators and signaling genes directly, and then regulate expressions of other genes through the ceRNA competition mechanism.

**Figure 4:**
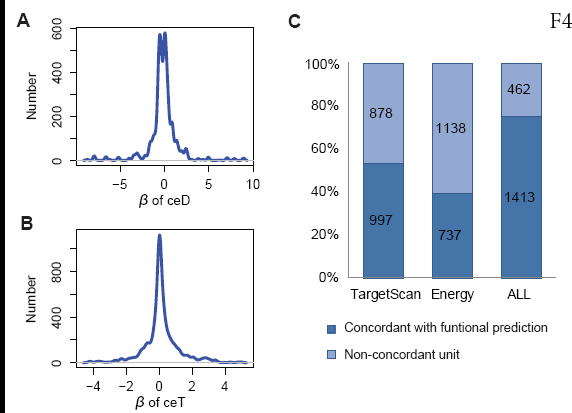
Figure 4: The functional properties of emsQTLs for EUR population. (A) The distribution of *β*_*d*_. (B) The distribution of *β*_*t*_. (C) The direction concordance between association and functional prediction for all emsQTLs.

The variant’s effect on miRNA-target interaction can be assessed by functional prediction algorithms that have been developed to estimate the change of binding affinity among different variant alleles^26; 42; 43^. To evaluate whether the direction of association (*β*_*d*_) is concordant with computational prediction on ceD through variant effect in *cis*, we calculated two scores, Δcontext+ score and ΔΔΔG, using TargetScan and an energy-based method^36^ for the 387 emsQTLs in EUR population. Intuitively, Δcontext+ score reflects the discrepancy of binding affinity and ΔΔΔG measures difference of combined interaction energy between alternative allele and reference allele. We found 53% and 39% of emsQTLs have *β*_*d*_ in consistent direction with Δcontext+ score and ΔΔΔG in functional prediction respectively (Figure 4C), and this increases to 75% if we consider the consistency with either Δcontext+ score or ΔΔΔG as independent validation (Figure 4C). These results suggest that majority of detected emsQTLs can be validated by functional prediction in their ceD locus.

The concordance of direction between statistical association and functional prediction can help us *in silico* prioritize the emsQTL candidates with potential causal evidence. From the 1,413 concordant variant-miRNA-ceD-ceT units for 250 emsQTLs, one can infer their molecular causality from both computational prediction and quantitative interpretation. Here, we use an example to illustrate how the emsQTL works. SNP rs1056984 is predicted to affect the seed binding between hsa-miR-296-5p and 3’UTR of *DIDO1*. TargetScan predicted binding under ancestral allele G (7mer-m8, context+ score: -0.236), but not under the derived allele A on this SNP. Further thermodynamic estimation confirmed that allele G has better binding affinity to miRNA (MFE of A allele: -26.7 kcal/mol; MFE of allele G: -30.5 kcal/mol). The simulated binding pattern also shows that allele G will enhance the binding stability by creating G:C match to position 8 of hsa-miR-296-5p (Figure 5A and 5B). Our model detected rs1056984 to be an emsQTL (*P*=4.76E-05) that controls the regulation among hsa-miR-296-5p, ENSG00000101191 (*DIDO1*) and ENSG00000185361 (*TNFAIP8L1*). The *β*_*d*_value of ceD *DIDO1* is -1.34 (Figure 5C), and the *β*_*t*_ value of ceT *TNFAIP8L1* is 0.20 (Figure 5D). The negative value of *β*_*d*_ further shows that rs1056984 is perhaps a gain-of-function mutation, which is consistent with functional prediction. This reversed relationship between coefficients indicates that genetic effect is driving the competing process of ceRNAs regulation. As the sequence of this allele changes from AA to AG to GG, the gradually enhanced sponge effect down-regulates *DIDO1* expression (ceD) and up-regulates *TNFAIP8L1* expression (ceT). When this locus is homozygous GG, we observed a significantly positive correlation (Cor = 0.29, *P* = 0.01) between *DIDO1* and *TNFAIP8L1*, further supporting the interaction between ceD and ceT through competition for hsa-miR-296-5p (Figure 5E).

**Figure 5:**
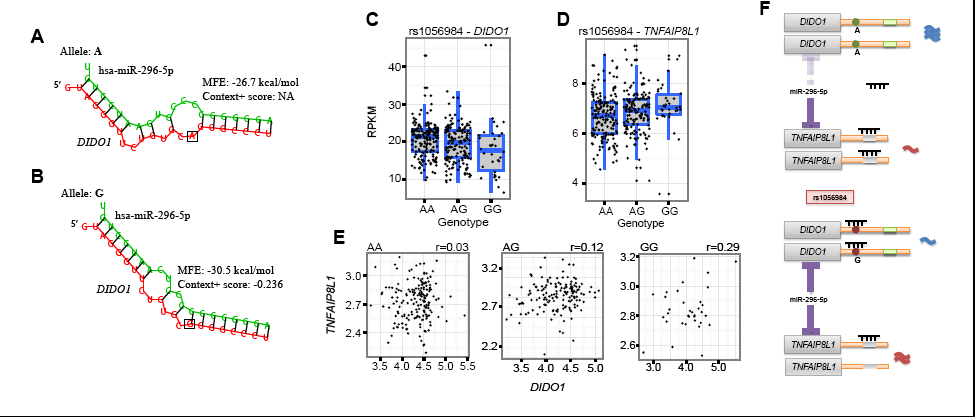
The genetic effect of rs1056984 in ceRNA regulation for EUR population. (A) Hybridization pattern between miR-296-5p and binding site of *DIDO1* on derived allele A. (B) Hybridization pattern between miR-296-5p and binding site of *DIDO1* on ancestral allele G. (C) Boxplot of gene expression of *DIDO1* on different genotype. (D) Boxplot of gene expression of *TNFAIP8L1* on different genotypes. (E) The gene expression correlation of *DIDO1* and *TNFAIP8L1* under different genotypes. (F) Schematic diagram for rs1056984 affecting ceRNA competition under different alleles, it impacts the expression of tumor suppressor and oncogene in a reciprocal and coordinate manner. MFE: minimum free energy.

When we searched biological functions for *DIDO1*, *TNFAIP8L1* and miR-296-5p, we found that this emsQTL may represent a new mechanism for miR-296-5p triggered carcinogenesis. miR-296 has been characterized as “angiomiR” which can regulate angiogenesis^44^. It is reported to have a specific role to promote tumor angiogenesis by targeting *HGS* mRNA and resulting in the overexpression of VEGF receptors in angiogenic endothelial cells^45-47^. MiR-296 may also contribute to carcinogenesis by dysregulating p53^48^. In this scenario, *DIDO1* gene is a tyrosine-phosphorylated putative transcription factor, previously thought to induce apoptosis and mitotic division^49; 50^, and might be a tumor suppressor gene^51^. In contrast, majority of publications reported *TNFAIP8L1* to be an antiapoptotic molecule and oncogene in developing many cancers^13; 52-54^. Here, we predicted emsQTL rs1056984 to affect the ceRNA regulation in switching the expression of tumor suppressor and oncogene under different genotypes. Specifically, efficient miRNA competition occurs in the ancestral allele G, however, the derived allele A of rs1056984 has a protective effect in maintaining tumor suppressor *DIDO1* expression and inhibiting oncogenic *TNFAIP8L1* expression by shifting miR-296-5p binding from *DIDO1* to *TNFAIP8L1* (Figure 5F). Although there is no diseases/traits associated evidences for rs1056984 at the current stage, we found that African population have lower derived allele frequency (DAF of YRI is 0.28, DAF of CEU is 0.65) in the 1000 Genomes project (Supplemental Figure S8). Further calculation on *F*_ST_ (0.24) between CEU and YRI indicates that positive selection may drive the evolution of this locus.

#### emsQTLs explain GWAS traits and diseases associated signals in miRNA binding sites

To investigate if emsQTL-affected gene expression changes contribute to human phenotypes, we connected emsQTLs in EUR population to GWAS trait/disease-associated SNPs (TASs), and found 8 of 387 ubiquitous emsQTLs to overlap with GWAS leading TASs (Table 1). The top mapped TAS rs7294, which locates in the 3’UTR region of *VKORC1*, has been frequently shown to be associated with warfarin maintenance dose in anticoagulant therapy^55-57^. Individuals with derived allele A produce less Vitamin K epoxide reductase than those with the G allele (“non-A haplotype”), thus the former need lower warfarin doses to inhibit the enzyme and produce an anticoagulant effect^58^. Our emsQTL analysis suggests that at the molecular level, allele A of rs7294 may promote the interaction among miR-147a and *VKORC1* target site (*β*_*d*_ = -1.12) to down-regulate *VKORC1* expression (Supplemental Figure S9A), consistent with the previously observed *VKORC1* expression in A allele individuals^58^. Also, the competition effect of this locus can help connect *VKORC1* to three significant ceTs *EIF2B5*, *LRRFIP1* and *RPTOR* (Supplemental Figure S9B-D) that are important in translational initiation or signaling pathway regulation.

**Table 1.** The emsQTLs that overlap with GWAS leading TASs

| Chr | Pos | SNP | Ref | Alt | emsQTL<br>P-value<sup>a</sup> | ceD | ceTs | miRNA | Effect<sup>b</sup> | GWAS<br>P-value<sup>c</sup> | GWAS Traits |
| --- | --- | --- | --- | --- | --- | --- | --- | --- | --- | --- | --- |
| 16 | 31102321 | rs7294 | C | T | 1.37E-06 | VKORC1 | EIF2B5, RPTOR,<br>LRRFIP1 | miR-147a | Gain | 1.40E-45 | Warfarin and acenocoumarol<br>maintenance dosage<sup>58, 78</sup> |
| 11 | 18388128 | rs4596 | G | C | 1.15E-05 | GTF2H1 | TSTD2, TMOD1,<br>TOB1, PPARGC1A,<br>KLF5 | miR-642a-5p | Loss | 2.17E-35 | Amyloid A Levels<sup>79</sup>, Colorectal<br>cancer<sup>80</sup> |
| 12 | 56863770 | rs2657880 | G | C | 1.87E-51 | SPRYD4 | XKR6, AC004985.2 | miR-3157-5p | Loss | 2.28E-31 | Serum metabolite levels<sup>81</sup>,<br>Lymphocyte counts<sup>82</sup>,<br>Metabolite levels<sup>83</sup> |
| 10 | 100176869 | rs701801 | C | T | 4.70E-13 | HPS1 | TRIOBP, KLHL30 | miR-491-5p | Loss | 1.34E-25 | Serum metabolism<sup>84</sup>, Endocrine<br>traits<sup>85</sup> |
| 19 | 10397238 | rs281437 | C | T | 0.00019 | ICAM1 | C16orf54,<br>FBXO41, SH2D4A | miR-3667-5p | Loss | 3.00E-10 | Soluble intercellular adhesion<br>molecule 1 level<sup>86</sup>, hepatic<br>fibrosis<sup>87</sup> |
| 16 | 67708897 | rs12449157 | A | G | 7.69E-06 | GFOD2 | COASY, LMAN2L,<br>VPS9D1 | miR-4792-5p | Gain | 2.00E-07 | Triglycerides<sup>88</sup>, Haemoglobin<br>level<sup>89</sup>,<br>Hypercholesterolemia<sup>90</sup> |
| 17 | 37921742 | rs907091 | C | T | 9.50E-05 | IKZF3 | RGL3, DDX11 | miR-330-5p | Gain | 3.38E-07 | Esophageal cancer<sup>91</sup>, Allergy<sup>92</sup><br>Primary biliary cirrhosis<sup>93</sup> |
| 8 | 11643915 | rs804292 | G | A | 0.00024 | NEIL2 | ZNF583 | miR-143-3p | Loss | 2.00E-06 | Alcohol/nicotine dependence<sup>94</sup> |

a: the best emsQTL P-value among all significant variant-miRNA-ceD-ceT units; b: the predicted function effect of alternative allele for miRNA-target interaction; c: the best GWAS P-value among all mapped GWAS traits

Since GWAS TAS may not be causal, we further scanned SNPs in high LD (r^2^ > 0.8) of GWAS-identified TASs. We identified 15.7% of 344 emsQTLs (SNPs only and not indels) to be strongly associated with 145 GWAS hits (Supplemental Table S5), an significant enrichment of emsQTLs in GWAS TASs comparing with background SNPs for both the miRNA seed binding site (*P* = 7.54E-28, hypergeometric test) and differentiated miRNA binding signals (*P* = 3.30E-4, hypergeometric test) (Supplemental Table S6). Most of these 145 GWAS index SNPs locate in introns or intergenic regions with poorly annotated functions (Supplemental Table S7). Therefore, our emsQTL analysis could potentially identify causal mechanisms underlying disease/trait SNPs. Interestingly, phenotypes associated with these 145 GWAS hits are mostly related to autoimmune diseases and blood cell traits (Supplemental Table S7), suggesting the effect of emsQTLs to be driven by cell type specificity of the lymphoblastoid cells in the Geuvadis data.

### Functional Effect of emsQTLs in ceRNA Regulatory Network

#### Prioritization of emsQTLs

To comprehensively evaluate the association between emsQTLs and ceRNA regulation, we rely not only on the statistical significance, but also on the magnitude of emsQTLs function on titrating miRNA availability and ceRNA-dependent gene expression changes. Several factors have been reported to influence ceRNA effectiveness, including miRNA and ceRNA expression level, the binding affinity of MRE, as well as the positive correlation between ceRNAs expression^3; 5; 59^. We attempted to prioritize the functional emsQTLs according to these factors. Since our regression model has already accounted for confounding factors from miRNA expression variation in the emsQTL calling step, we therefore only focused on ceRNA-related factors in the prioritization. We first calculated the degree of gene expression change on ceD and ceT in different genotypes, which can be measured by the sum of log|*β*_*d*_| and *log*|*β*_*t*_|. We further required consistent direction between *β*_*d*_ and the Δcontext+ functional prediction score from TargetScan. Finally, we asked for positive correlation (> 0.1) for ceD and ceT in the specific homozygous emsQTL genotypes, when ceD and ceT actively compete for miRNA binding. Based on aforementioned criteria, we successfully identified 239 variant-miRNA-ceD-ceT units for 93 unique emsQTLs with sufficient functional evidences (Supplemental Table S8). The top variant rs3208409 creates a miR-940-3p binding site in the 3’UTR of *HLA-DRB1* gene, which competes with *L3MBTL2* for miR-940-3p binding (*P* = 7.94E-27). The large effect of rs3208409 on the gene expression of ceD (*β*_*d*_ = -115.18) and ceT (*β*_*t*_ = 1.14), the consistent functional prediction (Δcontext+ score = -0.21), and the high correlation (Cor=0.38) between ceD and ceT in homozygote individuals provide robust evidences of the causality of this emsQTL (Supplemental Figure S10). We further overlapped this prioritized list with GWAS signals and identified 21 phenotype-associated variant-miRNA-ceD-ceT units (Supplemental Table S9).

#### Genetic effect on ceRNA regulatory network

The altered expression of individual genes might affect the expression of many other genes in the whole ceRNA regulatory network by the miRNA sponge mechanism^60; 61^. From the 1,875 significant variant-miRNA-ceD-ceT units our model identified, we constructed the global ceRNA regulatory network under the control of 387 independent emsQTLs in the EUR population (Supplemental Figure S11). We also generated the network for 21 phenotype-associated variant-miRNA-ceD-ceT units (Supplemental Figure S12). Majority of ceDs can be associated with more than one ceTs by single genetic effect. For example, rs11540855 on ceD *ABHD8* could influence the expression of two ceTs *AXIN1* and *RPRM* through competing for binding to miR-4707-3p (Figure 6A), and the expression of *ABHD8* and its two ceTs are positively correlated under the active genotype GG (Figure 6B-D). Interestingly, emsQTL rs11540855 located in the 3’ UTR of *ABHD8* on 19p13 has been recently reported to have top significant association with breast cancer risk (GWAS *P* = 1.65E-09) after genotype imputation^62; 63^. In addition, *AXIN1* and *RPRM* were recently reported as tumor suppressors in breast cancer development^64^, and miR-4707-3p is highly expressed in breast cancer^65^. These evidences suggest that emsQTL rs11540855 might influence breast cancer developments by regulating tumor suppressors *AXIN1* and *RPRM* through the ceRNA pathway. In addition to the aforementioned regulatory relationship, one ceD can also be regulated by multiple miRNAs, and a single miRNA can regulate multiple ceDs and ceTs through different emsQTLs. These interactions highlight the complexity of genetic effect on ceRNA regulatory network.

**Figure 6:**
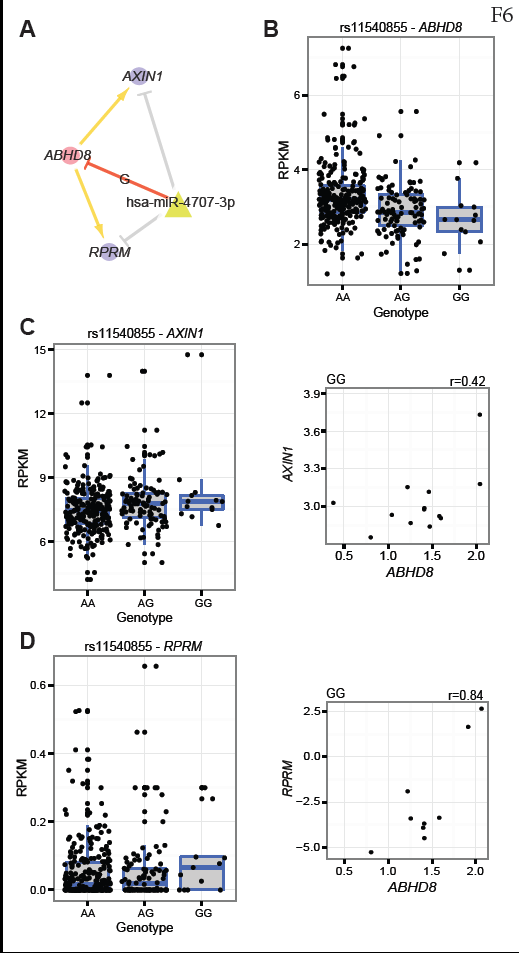
The genetic effect of rs11540855 in ceRNA regulation for EUR population. (A) Small ceRNA regulatory network driven by rs11540855, red circle: ceDs; yellow triangle: miRNA; bule circle: ceTs; red suppression line: the miRNA-ceD regulation, G for gain-of-function mutation; gray suppression line: the miRNA-ceT regulation; orange arrow: ceD activate ceTs in gain-of-function situation (βd < 0 and βt > 0). (B) Boxplot of gene expression of *ABHD8* on different genotypes. (C) Boxplot of gene expression of *AXIN1* on different genotypes and the correlation with *ABHD8* on genotype GG. (C) Boxplot of gene expression of *RPRM* on different genotypes and the correlation with *ABHD8* on genotype GG.

## Discussion

In this study, we, for the first time, integrated the 1000 Genomes genotype and Geuvadis RNA sequencing data to investigate the effect of human genetic variations on ceRNA regulation. Using a multivariate linear model, we successfully identified hundreds of emsQTLs and related ceDs/ceTs at the genome-wide level. We found that recent natural selection is shaping many emsQTLs in different human populations. Functional analysis of these genetic variants indicated that most of emsQTLs are functionally relevant to important biological processes and are significantly enriched in GWAS risk loci. Furthermore, we prioritized these loci with their associated ceRNAs according to different criteria and evaluated their collective effect on the ceRNA regulatory network. Our study provides a novel angle to interpret genetic effect in post-transcriptional gene regulation.

Although our regression model already considered many candidate confounding factors, such as miRNA expression level, ceRNA expression variability, as well as population genetic structure, there may still be missing factors that impact the performance and statistical power in the emsQTL detection. One potential limitation is that we only treated each pair of ceD and ceT as an independent test unit in the local miRNA-centered regulatory network instead of modeling the whole ceRNA regulatory network. Recent studies have shown that a small perturbation of ceRNA expression usually shifts the equilibrium of ceRNA regulatory network especially when concentrations of miRNAs and targets are comparable^66^. The cascade effect from miRNA redistribution and ceRNA competition in the global level^8; 67^ requires a complete and complex mathematical model to accurately describe full responses of the whole network. Another limitation of our study is that current computational predictions of miRNA binding sites still have inadequate performances^68^. To balance sensitivity and specificity of miRNA target prediction, we chose to use a strict context+ score threshold from TargetScan predictions, instead of the union or intersection of multiple miRNA-target prediction algorithms such as TargetScan, PITA^36^, miRanda^69^, etc. Therefore, it is likely that our emsQTLs detection missed some causal variants not predicted by TargetScan. Future experiments such as CLIP-seq, if done on individuals, could better capture miRNA-target interactions and improve our emsQTLs inference.

Using genetic and transcriptomic data from different populations, we found many population-specific emsQTLs and identified putative loci undergoing recent positive selection. These results represent a useful supplement to studies of recent natural selection of human miRNA targets^70-72^and significantly extent functional categories for positively selected loci^73^. Currently, available transcriptome profiles of five subpopulation from Geuvadis, of which majority are from European populations, prevent the inference of positive selection signals in other human races such as Asian and Native American populations. The recent human Genotype-Tissue Expression (GTEx) project has produced large-scale transcriptome profiles in multiple tissues of hundreds of donors^74^, which provides new opportunities to study tissue-specific associations between genetic variations and ceRNA regulation.

Since many genes contain multiple binding sites of the same miRNA, some might suspect single MRE perturbations to have small effects on ceRNA expression and the downstream miRNA regulatory network. This is not surprising, considering that most QTLs usually also only account for a small fraction of the total genetic heritability in the population. However, although many QTLs individually exert relatively small effects, together they might contribute to a significant complex trait^75^. Theoretical simulations and quantitative experiments have demonstrated that some perturbations on individual miRNA binding site can indeed affect the entire ceRNA regulatory network^8; 76; 77^. Our emsQTL analyses on human populations suggest that DNA polymorphisms affecting ceRNA regulation is a widespread phenomenon in the human evolution and contribute significantly to complex traits.

## Description of Supplemental Data

Supplemental Data include twelve figures and nine tables.

## Acknowledgement

We are very grateful to Dr. Graham McVicker for his discussions and suggestions. The project was supported by funds from Y S and Christabel Lung Postgraduate Scholarship (MJL), Research Grants Council, Hong Kong SAR, China 17121414M (JWW), The National Institute of Health U41 HG7000 (XSL) and The National Institute of Health R01 GM113242-01 (JSL).

